# Human dimension of Holocene wildfire dynamics in boreal eastern Siberia

**DOI:** 10.1101/2025.03.14.643308

**Authors:** Ramesh Glückler, Elisabeth Dietze, Andrei A. Andreev, Stefan Kruse, Evgenii S. Zakharov, Izabella A. Baisheva, Amelie Stieg, Shiro Tsuyuzaki, Luidmila A. Pestryakova, Ulrike Herzschuh

**Affiliations:** Polar Terrestrial Environmental Systems, Alfred Wegener Institute Helmholtz Centre for Polar and Marine Research, Potsdam, Germany; Institute for Environmental Science and Geography, University of Potsdam, Potsdam, Germany; Faculty of Environmental Earth Science, Hokkaido University, Sapporo, Japan; Institute of Geography, Georg August University of Göttingen, Göttingen, Germany; Institute of Natural Sciences, North-Eastern Federal University of Yakutsk, Yakutsk, Russia; Institute for Biochemistry and Biology, University of Potsdam, Potsdam, Germany

## Abstract

Severe wildfire seasons in the Republic of Sakha (Yakutia) raise questions regarding long-term fire dynamics and their drivers. However, data on long-term fire history remains scarce across eastern Siberia. We present the first composite of reconstructed wildfire dynamics in Yakutia throughout the Holocene, based on eight newly contributed records of macroscopic charcoal in lake sediments in combination with published data. Increased biomass burning occurred in the Early Holocene, c. 10,000 years BP, before shifting to lower levels at c. 6000 years BP. Independent simulations of climate-driven burned area in an individual-based forest model reproduce this reconstructed Holocene trend (*r*_s_ = 0.85), but the correlation on centennial timescales turns negative in the Late Holocene (*r*_s_ = -0.70). This mismatch suggests that climate alone cannot explain Late Holocene wildfire dynamics. We propose that a human dimension needs to be considered. By example of the settlement of the pastoralist Sakha people c. 800 years BP, we show that implementing reduced fuel availability from Indigenous land management in the forest model leads to improved centennial-scale correlations (*r*_s_ = 0.96). This study highlights the need for a better understanding of the poorly reported human dimension of past fire dynamics in eastern Siberia.

## Introduction

Wildfires in boreal forests are becoming more extreme (Cunningham et al., 2024), with eastern Siberia as one of the most affected regions (Jones et al., 2022). The fire regime intensification here occurs in a unique and climate-sensitive ecosystem of deciduous larch forests and continuous permafrost (Kharuk et al., 2021). Annual burned area and fire intensity in the Republic of Sakha (Yakutia), referred to as the coldest permanently inhabited region on Earth (Li, 2016), have seen pronounced increases over recent decades (Li et al., 2024). Fire brigades and voluntary firefighters across Yakutia, challenged by a lack of funding and restrictive policies (Narita et al., 2021; Canosa et al., 2023), were confronted with unusually intense crown fires engulfing whole tree stands. As a consequence, several settlements were endangered, infrastructure was blocked or destroyed, and exposure to hazardous smoke was widespread (Tomshin and Solovyev, 2022; Vinokurova et al., 2022). It is predicted that fire regimes will continue to intensify under a steadily warming climate (Jones et al., 2022; Burton et al., 2024), with unknown implications for ecosystem stability (Furyaev et al., 2001; Shestakova et al., 2024). In addition, the extent to which humans may have affected past fire regimes is unknown and remains difficult to assess (Marlon et al., 2016). Systematic satellite-based fire observations over the last few decades are a valuable source of data; however, they do not permit a direct evaluation of centennial to millennial-scale trends. An evaluation of past natural and human drivers behind changing wildfire dynamics requires information for more than the last few decades.

The long-term fire history is unknown across most regions of the vast larch forests of eastern Siberia, covering c. 2.6 million km^2^ and representing almost 40% of the forested area of Russia (Abaimov, 2010). This is due to a lack of paleoecological data on past wildfire dynamics (Marlon et al., 2016). Only a few studies have investigated centennial-to millennial-scale fire dynamics in boreal Yakutia (Pupysheva and Blyakharchuk, 2023). Macroscopic charcoal particles (>100 µm) in lake sediments can be leveraged as a paleoecological proxy to reconstruct trends of biomass burning in the vicinity of a lake (Whitlock and Larsen, 2001). Katamura et al. (2009ab) contributed charcoal records near Yakutsk, spanning the Holocene (Lake Sugun, Lake Chai-ku, Maralay Alaas; Fig. 1). Based on their data, they suggest a stable surface fire regime throughout the Holocene. In contrast, Glückler et al. (2022) find variable Holocene fire activity and the establishment of the modern surface fire regime c. 4500 years BP (Before Present; i.e., before calendar year 1950) from a charcoal record in western Central Yakutia (Lake Satagay; Fig. 1). Besides the scarce data availability limiting the understanding of Holocene fire history, underlying drivers have not, so far, been systematically investigated.

**Fig. 1:**
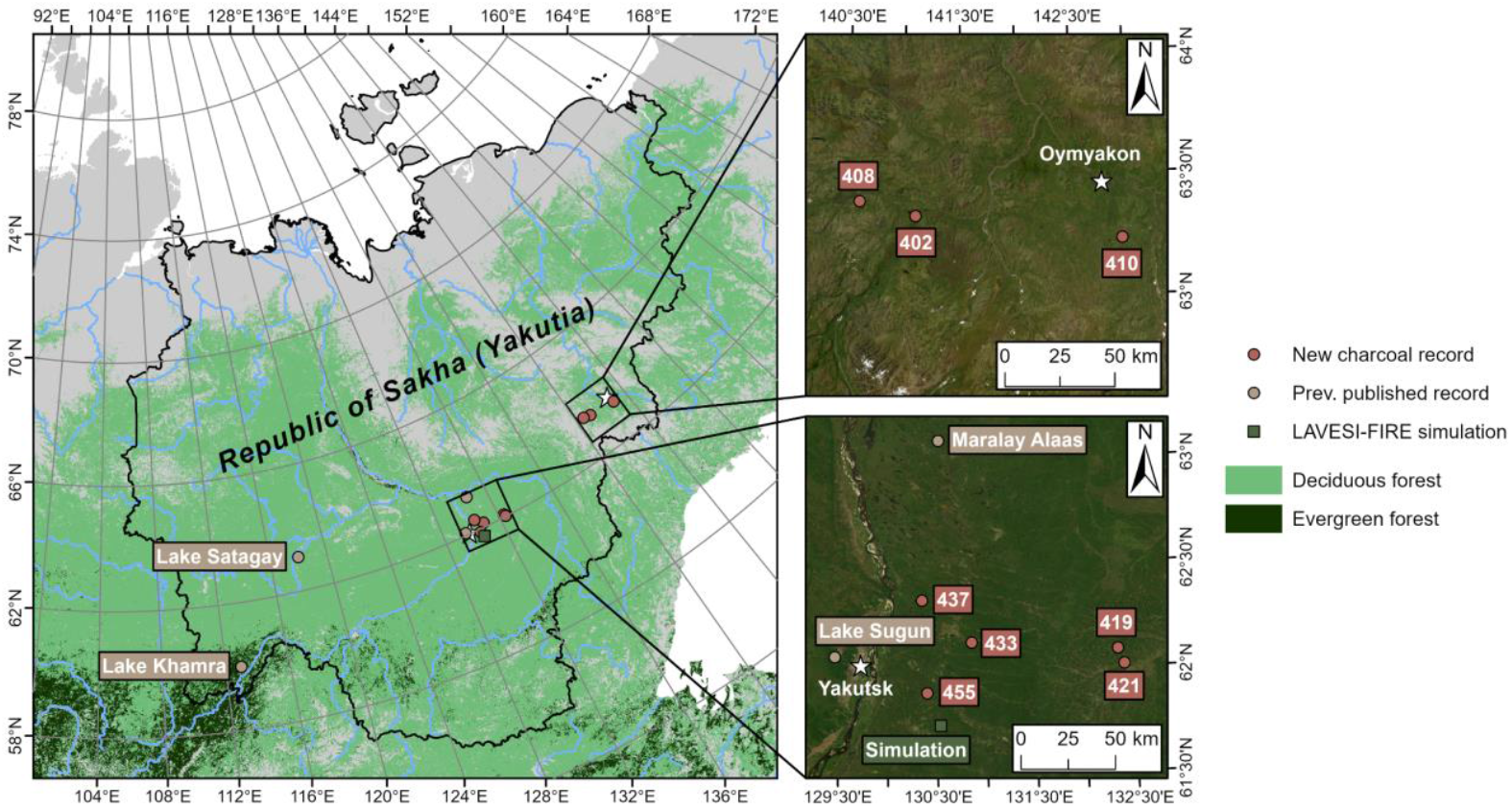
Locations of lake-sedimentary charcoal records and simulations in LAVESI-FIRE. Blue lines mark major river centerlines from Natural Earth. Boreal forest extent from ESA Land Cover CCI. Coordinate systems used: WGS 1984 EPSG Russia Polar Stereographic (left), UTM 54N (upper right), UTM 52N (lower right). “World imagery” basemap sources: Esri, DigitalGlobe, GeoEye, i-cubed, USDA FSA, USGS, AEX, Getmapping, Aerogrid, IGN, IGP, swisstopo, and the GIS User Community.

Climate constitutes an important natural driver of long-term fire regime changes. Temperature and relative humidity are key factors influencing fire weather (Jain et al., 2022). Climatic variables can therefore affect the number of ignitions, fire extent, and fire severity (Sayedi et al., 2024). However, the impact of fire weather on fire regimes varies depending on other factors, such as fuel availability, and can therefore be heterogeneous in space and time. At Lake Satagay, reconstructed periods with high amounts of biomass burning were suggested to correspond to known climatic phases, such as the Holocene Climatic Optimum (Glückler et al., 2022). A closer analysis of climate as a driver of Holocene wildfire dynamics was hampered by the low availability of paleoclimatological data from the region. As a result, the influence of past climatic changes on fire dynamics in Yakutia is poorly understood.

Humans were found to act as drivers of fire regimes for many millennia in various regions of the world (Girardin et al., 2024; Bird et al., 2024). However, the extent to which people in boreal Yakutia influenced past wildfire dynamics is unknown. Hominins lived across southern Siberia at least c. 800,000 years ago (Kuzmin et al., 2022), and humans from nomadic hunter-gatherer cultures roamed eastern Siberia since more than 40,000 years ago (Pitulko et al., 2016; Sikora et al., 2019). An increased frequency of occupation periods across eastern Siberia has been recorded since the Mid-Holocene, and Central Yakutia may have been a prominent station along south-to-north migration routes (Kuzmin, 2023). It is not known whether these activities influenced fire regimes. The arrival of the pastoralist Sakha people in Central Yakutia around 1200 CE (Common Era) resulted in a cultural shift towards dispersed semi-nomadic and sedentary livelihoods based on horse and cattle breeding (Fedorova et al., 2013). Many traditional land management practices, such as hay making, shrub removal, and cultural burning (Okladnikov, 1970; Crate, 2021), likely had complex impacts on the environment. Whether the probable consequence of reduced fuel loads may have affected fire regimes has not yet been tested. In a high-resolution charcoal record, covering the last two millennia in southwest Yakutia (Lake Khamra; Fig. 1), Glückler et al. (2021) found high wildfire activity around 900 CE and low activity until c. 1850 CE. Due to a stable vegetation composition, it was speculated that changes in fire regime were initially driven by climatic changes and increasingly influenced by human land management only during the most recent centuries. However, these speculations were limited by the evidence being based on a single site and the lack of a specific method to test for changing natural or human drivers.

Regional trends of biomass burning during wildfires can be derived from the more locally-influenced macroscopic charcoal records by synthesizing a composite curve out of multiple sites (Power et al., 2008; Marlon et al., 2013). This approach, necessitating a higher availability of charcoal records, enables an analysis of past wildfire dynamics beyond the local scale of each individual record. While composited charcoal records provide data on past wildfire dynamics, global climate simulations in earth system models can be used to determine causal relationships with climate data and assess potential climatic drivers (Dallmeyer et al., 2022). Such climate data can also be used to directly simulate wildfires in forest models for eastern Siberia. Modeling frameworks can therefore provide a novel opportunity to test for potential impacts of past climate and human land management practices on wildfire dynamics. Of the few forest models localized specifically to capture fine-scale larch forest dynamics (e.g., SEIB-DGVM, Sato et al., 2010; UVAFME, Shuman et al., 2017), only the individual-based, spatially explicit LAVESI-FIRE was previously used to validate multi-millennial trends of larch forest development and wildfire disturbance in an initial comparison with paleoecological data from a charcoal record (Glückler et al., 2024).

In this study, we aim to elucidate Holocene wildfire history in Yakutia and to disentangle natural from potential human drivers. We contribute eight new Holocene records of macroscopic charcoal in lake sediments and simulate climate-driven wildfires in a localized forest model. By creating a composite curve of Holocene charcoal accumulation, we analyze long-term wildfire dynamics. We then simulate climate-driven burned area to estimate the role of climate as a driver behind reconstructed fire dynamics, and to test the hypothesis of reduced fuel availability impacting fire regimes in the Late Holocene. Results reveal that a reconstructed severe fire regime during the Early to Mid-Holocene corresponds well to the estimated impact of past climatic trends. While climate may be able to explain longer multi-millennial trends of biomass burning, shorter multi-centennial trends since the Mid-Holocene may rather result from human activity. By example of the Sakha people, we discuss how traditional land management practices may have influenced fire regimes long before industrialization.

## Results

### Reconstructed Holocene wildfire dynamics

Eight newly contributed records of the accumulation of macroscopic charcoal particles in lake sediments from the Lena-Amga interfluve in the Central Yakutian Lowland, the southern Verkhoyansk Mountains, and the Oymyakon Highlands highly increase Holocene paleofire data availability in Yakutia (Fig. 1; Supplement 5). The new records cover the last 7200 years or parts of this period. The average charcoal accumulation rate for all newly contributed charcoal records is 0.33 ± 0.41 particles cm^-2^ yr^-1^ (mean ± standard deviation), ranging for individual records from a minimum of 0.04 ± 0.04 particles cm^-2^ yr^-1^ (Lake 402, Lake 408) to a maximum of 1.27 ± 1.08 particles cm^-2^ yr^-1^ (Lake 421).

We obtained the regional trend of charcoal accumulation throughout most of the Holocene (10,930 to -70 years BP; Before Present) by compositing the eight new records in combination with four existing records from the region in the Global Paleofire Database (formerly known as Global Charcoal Database; Power et al., 2010; Marlon et al., 2016; Fig. 2). With an average temporal resolution of 82 ± 56 years per sample across all sediment cores, the composite curve highlights both millennial and centennial trends.

**Fig. 2:**
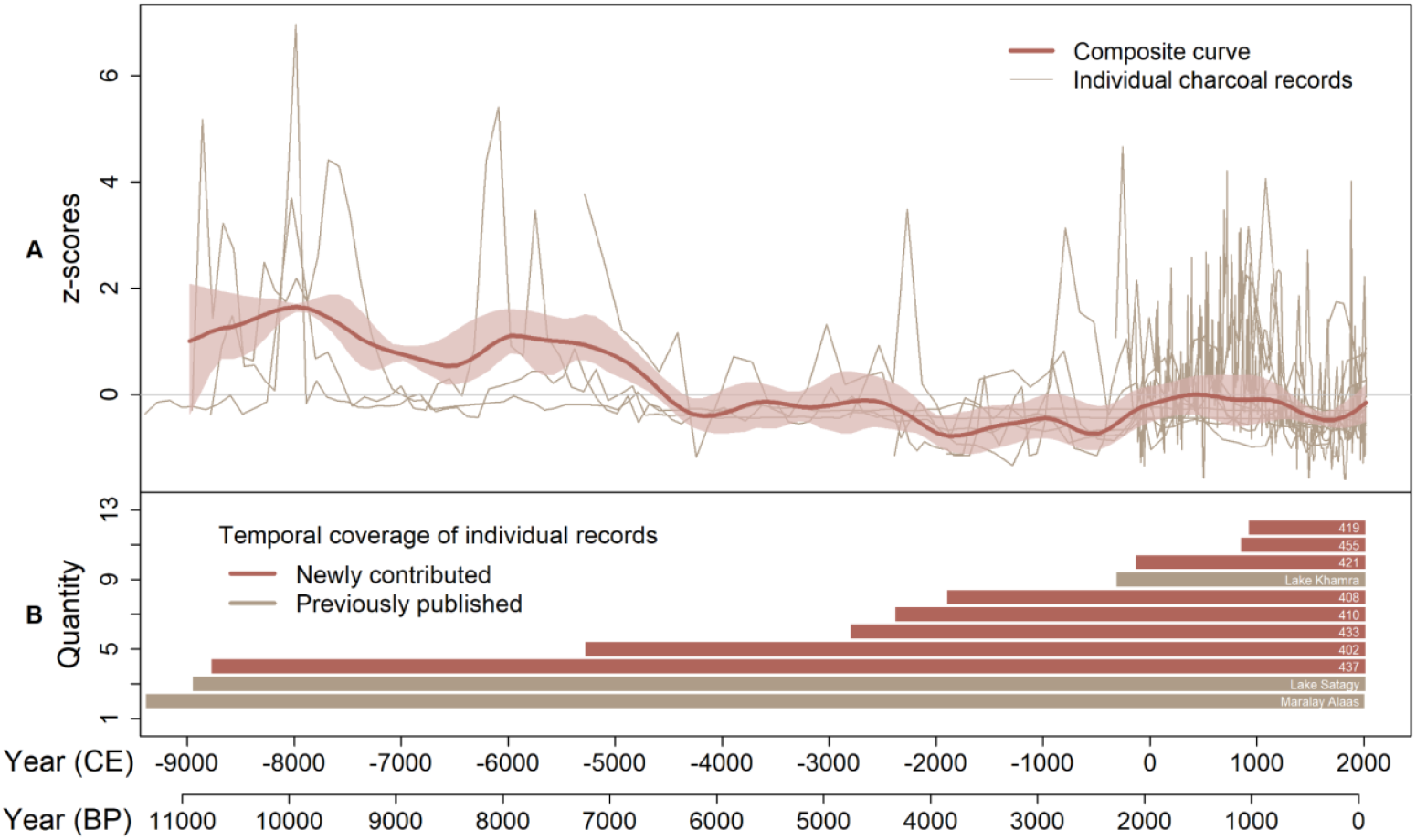
Charcoal accumulation rate throughout the Holocene indicates a trend from higher (Early Holocene) to lower (Late Holocene) levels of biomass burning. A: Composite curve of Yakutian charcoal records, standardized. Shaded area represents the 95% confidence interval. B: Temporal coverage of individual records included in the composite curve.

Charcoal accumulation during the Early Holocene and early Mid-Holocene was high, with two maxima at c. 10,000 and c. 8000 to 7000 years BP. In the Mid-Holocene, our results indicate a shift to a lower level after 6000 years BP. Charcoal accumulation continues to vary at this lower level throughout the Late Holocene, with record-wide minima at 4000 and 2500 years BP. The last two millennia are characterized by increasing values after 2500 years BP, followed by a pronounced decrease around 1200 CE. The most recent centuries see an increase in charcoal accumulation again, although remaining below levels recorded both in the Early Holocene and before 1200 CE. Confidence intervals of the composite curve suggest highest agreements between the different contributing charcoal records for the Early Holocene maximum (10,000 years BP) and the subsequent Mid-Holocene decrease (6000 years BP). Notably, high agreement is also suggested for the most recent decrease of charcoal accumulation (800 years BP or 1200 CE), where 11 out of 12 charcoal records are represented.

### Simulated climate-driven wildfire activity

We used the individual-based, spatially explicit forest model LAVESI-FIRE (Glückler et al., 2024) to simulate climate-driven burned area in Central Yakutia throughout the Holocene (simulation location is indicated in Fig. 1). Trends of simulated burned area compare well to the charcoal-based, reconstructed composite curve on a multi-millennial scale throughout the Holocene (Fig. 3B; Spearman rank correlation coefficient *r*_s_ = 0.85).

**Fig. 3:**
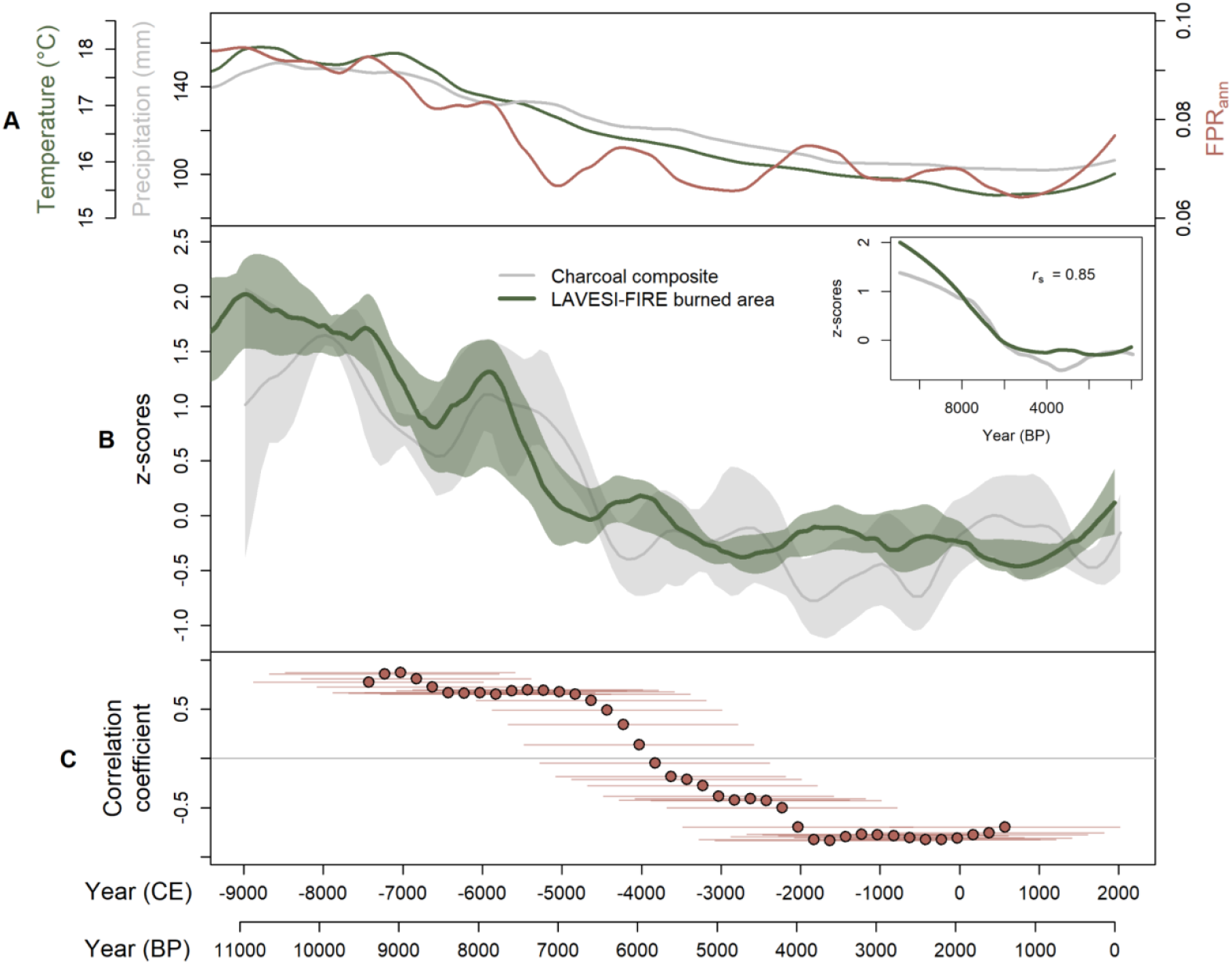
LAVESI-FIRE simulated burned area in comparison to the charcoal composite curve. On a longer multi-millennial scale both timeseries are highly correlated, but on a shorter multi-centennial scale their correlation turns negative in the Mid-Holocene. A: Climate forcing data for LAVESI-FIRE simulations from MPI-ESM-CR (Kapsch et al., 2022; smoothed annual June, July, and August mean temperature and precipitation sum) and the derived annual fire probability rating (FPR_ann_; Glückler et al., 2024). B: Comparison between annually simulated burned area and the charcoal composite curve. The green line represents the median of ten smoothed simulation repeats, while the transparent band marks the interquartile range. Inlay shows correlation between both timeseries on a multi-millennial scale. C: Moving window correlation coefficient (*r*_s_) between simulated burned area (median) and the charcoal composite curve (mean) on a multi-centennial scale. Points mark the center of each analyzed window, with horizontal lines indicating the full corresponding window extent.

A comparative analysis on a shorter multi-centennial scale (Fig. 3C) indicates that simulated burned area reproduces the high-agreement maximum of reconstructed charcoal accumulation at c. 10,000 years BP and the subsequent decrease before 6000 years BP. However, during the Mid-Holocene there is a shift from significant positive correlations (c. 11,000 to 5000 years BP) towards significant negative correlations (c. 5000 years BP to present). In the Late Holocene, reconstructed trends of biomass burning are not captured by climate-driven simulated burned area on these shorter timescales, including the last millennium (Fig. 4; *r*_s_ = -0.70). Artificially reducing the model’s fuel availability after 1200 CE, as a suggested net effect of human pastoralist activity, leads to an improved fit of simulated burned area to reconstructed trends of charcoal accumulation during this last millennium (Fig. 4; *r*_s_ = 0.96).

**Fig. 4:**
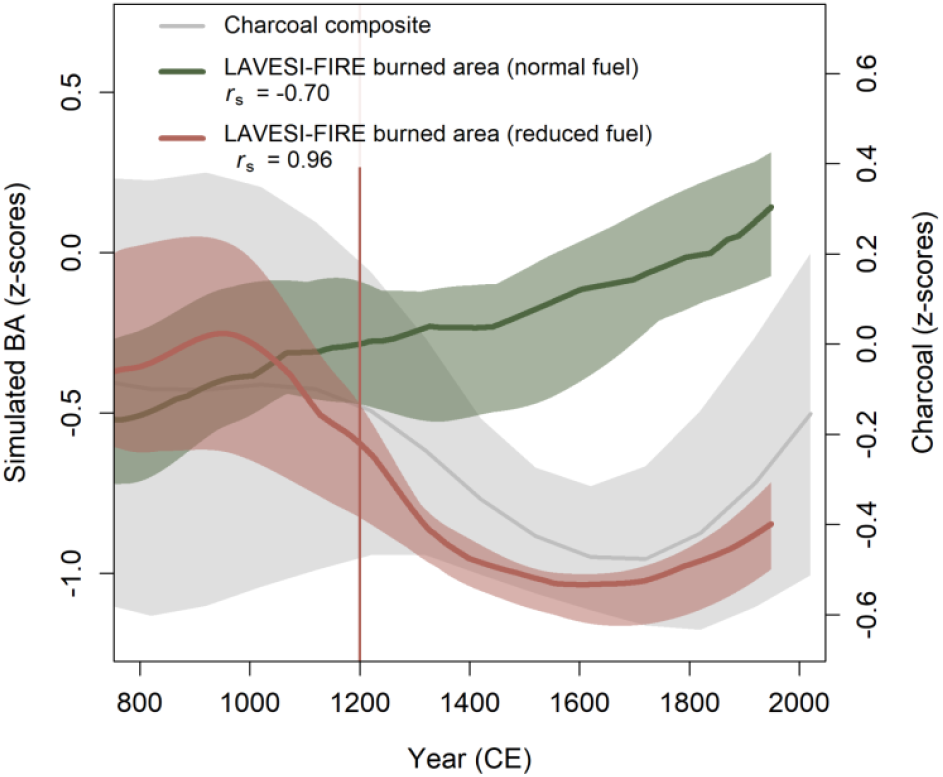
LAVESI-FIRE simulated burned area with (red) and without (green) artificial fuel reduction after 1200 CE, in comparison to the charcoal composite curve. Reducing the model’s fuel availability, as a suggested consequence of the settlement of the pastoralist Sakha people, improves the correlation between simulated and reconstructed data. Colored lines represent the median of ten simulation repeats per scenario, while the transparent bands mark the interquartile range. The vertical line at 1200 CE marks the beginning of the artificial fuel reduction.

## Discussion

### Holocene wildfire dynamics in eastern Siberia

Our composite curve in Yakutia indicates high regional wildfire activity in the Early to Mid-Holocene (c. 11,000 to 7000 years BP). This is followed by a shift to a lower level until present. This finding represents a deviation from reconstructed global and northern hemispheric biomass burning trends, where a long-term increase of biomass burning was reported since the beginning of the Holocene (Marlon et al., 2016). When assessed regionally, charcoal record composites reveal contrasting Holocene trends (Marlon et al., 2013). Our new data on long-term wildfire activity in Yakutia, as the first regional composite in eastern Siberia, differs from a regional-scale composite of charcoal records from interior Alaska, where gradually increasing biomass burning throughout the past 10,000 years was reported (Kelly et al., 2013). Power et al. (2008), on the other hand, found a maximum of biomass burning in the Early Holocene in eastern North America. Our regional composite from eastern Siberia fills an important data gap in the distribution of boreal paleofire studies (Marlon et al., 2016) and demonstrates that global or hemispheric composites are not necessarily representative of regional wildfire history in Yakutia, an especially fire-prone region (Kirillina et al., 2020).

The Holocene-scale trend of biomass burning is overlain by variation on a multi-centennial scale (Fig. 3B). Few charcoal records currently cover the Early to Mid-Holocene in Yakutia (Fig. 2B) and eastern Siberia as a whole. During this time, we find maxima of biomass burning at 10,000 and 8000 years BP, which is in agreement with maxima reported from macro- and microscopic charcoal records near Lake Baikal (Barhoumi et al., 2021; Krikunova et al., 2024).

These studies furthermore indicate rapid shifts to lower levels of biomass burning in the Mid-Holocene, although timing differs between lakes and is recorded about one millennium after the pronounced decrease around 6000 years BP we find in Yakutia. In the Mid-to Late Holocene, where more charcoal records are available, we record short-term minima of biomass burning at c. 4000 and 2500 years BP, followed by a pronounced decrease around 1200 CE. Similar trends have been found in other composites from the Northern Hemisphere, albeit at differing times (Marlon et al., 2016). Marlon et al. (2009) reported a decrease in burning from 1400 to 1750 CE. In south-central Canada, a marked decrease of biomass burning to low levels occurred around 2000 years BP (Girardin et al., 2024).

### Climate as major driver of long-term fire trend

We use a comparison between reconstructed and simulated wildfire activity to estimate the extent to which climate acted as a driver of past wildfire dynamics. Our model specifically represents climate-driven changes of the simulated burned area, since climate (expressed by monthly temperature and precipitation) is the main driver of fire occurrence, which is then only mediated by factors such as fuel availability and topography-derived surface moisture availability (Glückler et al., 2024). The simulation outcome tends to react more sensitively to temperature than precipitation, fitting to the largely temperature-limited environment (Glückler et al., 2024). This results in the simulated burned area being highly correlated to the climate-derived fire probability (Fig. 3A, B).

Climatic changes can explain our reconstructed wildfire dynamics on a longer multi-millennial scale throughout the Holocene. This is indicated by a high positive correlation between our reconstructed wildfire activity and simulated burned area, which is mainly driven by the climate (Fig. 3B; *r*_s_ = 0.85). This finding reinforces a previous assertion that long-term wildfire activity is strongly driven by climatic changes (Marlon et al., 2009), and we now show this to be the case in eastern Siberia as well. Following the Pleistocene-Holocene transition, a strongly warming climate resulted in the Holocene Climate Optimum (Tarasov et al., 2007), which has been suggested to facilitate high amounts of biomass burning in previous local studies (Bezrukova et al., 2011; Glückler et al., 2022). These climatic controls likely explain the high correlation between simulated and reconstructed data of this study on a longer, multi-millennial scale, characterized by wildfire activity shifting from high to low levels over the course of the Holocene.

Trends of reconstructed Holocene wildfire activity in Yakutia correspond to known climatic changes from independent climate reconstructions. Temperature estimates derived from regional pollen records in Central Yakutia confirm the multi-millennial trend of an Early Holocene increase shifting to a decrease in the Mid-Holocene, with a Holocene maximum around 8000 to 9000 years BP (Herzschuh et al., 2023). Lakes in Central Yakutia and the Yana Highlands recorded high productivity from increased organic carbon accumulation between c. 11,500 to 9000 years BP, likely facilitated by warm climatic conditions following peak insolation at the beginning of the Holocene (Baumer et al., 2020; Hughes-Allen et al., 2021). A cooling trend towards the Late Holocene is also reflected by oxygen isotopes in diatom silica of lakes across northern Eurasia (Meister et al., 2024), as well as in ice wedges in Central Yakutia (Popp et al., 2006).

On a shorter multi-centennial scale, however, trends of reconstructed and simulated wildfire activity are positively correlated only in the Early to Mid-Holocene (until c. 5000 years BP; Fig. 3B). During this time, our results show a period of reduced biomass burning around 8200 years BP. This coincides with a well-known global event following freshwater outbursts from the melting Laurentide ice shield, leading to a wide-spread decrease in temperature (Thomas et al., 2007; Parker and Harrison, 2022). The return to higher wildfire activity from c. 8000 to 7000 years BP corresponds to a warm period during the late stages of the Holocene Climate Optimum, which has been linked to widespread thermokarst initiation (Ulrich et al., 2017). A paleohydrological record from southern Siberia indicates unstable climatic conditions during the Early Holocene (Harding et al., 2020), which may explain the highly variable charcoal accumulation rates during that time (Fig. 2A). During the Late Holocene, climatic shifts of a lower amplitude, such as the Medieval Warm Period (c. 950 to 1250 CE) or the Little Ice Age (c. 1400 to 1750 CE; Mann et al., 2009), may have impacted wildfire activity. However, our climate-driven simulations are negatively correlated to reconstructed wildfire activity on the shorter multi-centennial scale (Fig. 3C), indicating that climate may not be mainly responsible for the reconstructed pattern of biomass burning during this time.

Vegetation composition, a key control of fire regimes by affecting fuel type and availability (Sayedi et al., 2024), did not change strongly throughout the Holocene and is thus unlikely to be a key driver of multi-centennial fire regime variability in the Late Holocene (Glückler et al., 2022). Reconstructions based on palynological data and sedimentary ancient DNA analyses show how Early Holocene open woodlands of Yakutia, populated mainly by larch (*Larix*) and birch (*Betula*), became gradually denser and mixed with only a few other species during the Holocene (Velichko et al., 1997; Müller et al., 2009; Andreev et al., 2022; Baisheva et al., 2024). Larch remained the main forest-forming tree ever since forest establishment began in tundra-steppe environments following the Last Glacial Maximum (Schulte et al., 2022). Vegetation composition may, instead, have reacted to variability in the fire regime. This is demonstrated by an increased abundance of birch, which commonly establishes on post-fire disturbed areas, during the fire-prone Early Holocene (Abaimov and Sofronov, 1996; Glückler et al., 2022). Reduced wildfire activity in the Mid-to Late Holocene, on the other hand, may have enabled evergreen trees such as the fire-avoiding spruce (*Picea*) to establish, albeit only in the southwest of Yakutia (Wirth, 2005; Glückler et al., 2021). Pine (*Pinus*) invaded Central Yakutia only in the Mid-Holocene, and remained limited to patches of sandy terrain such as along the Lena River banks (Andreev and Tarasov, 2013; Kharuk et al., 2021).

### Humans modifying centennial-scale fire trends since the Mid-Holocene

Neither vegetation composition nor climate appear to be the main driver behind the multi-centennial variability of reconstructed wildfire activity during the Late Holocene. Our results indicate a potential change in drivers around 5000 years BP, when the correlation of reconstructed biomass burning and simulated burned area first decreases to zero and then turns negative for the remaining period of the Late Holocene (Fig. 3C). This timing coincides with an increase in human activities in eastern Siberia.

Impacts of nomadic hunter-gatherers, who lived in eastern Siberia since before the Holocene (Pitulko et al., 2016), may have been limited in spatial scale and non-persistent during Mesolithic to Neolithic times (c. 12,000 to 5000 years BP; Fig. 5). Lightning-fire-prone environments (which includes boreal eastern Siberia) have been found to predict the use of fire by hunter-gatherer communities (Coughlan et al., 2018). In such environments, anthropogenic burning by small communities has the potential to decrease the severity of naturally occurring wildfires (Coughlan et al., 2018). It remains unclear to what degree this may apply to hunter-gatherers in eastern Siberia. Our results do not provide evidence for impacts of nomadic communities focused on hunting small mammals, fishing, and gathering plants (Kuzmin et al., 2022) on wildfire dynamics during this time. Therefore, we assume that any imprint in centennial to millennial-scale trends of wildfire activity beyond natural trends may have been limited (Roos et al., 2019; Ryabogina et al., 2024).

**Fig. 5:**
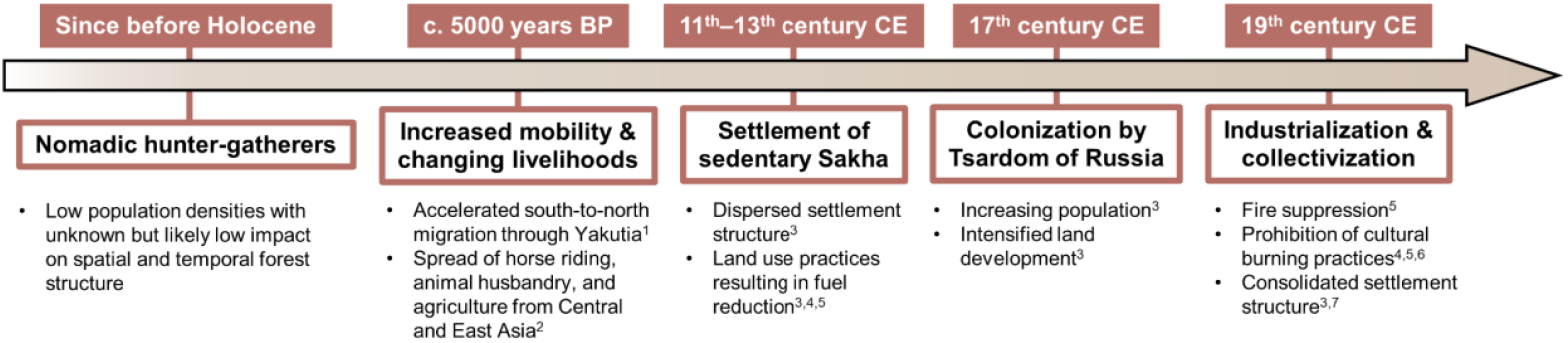
Schematic timeline of general important cultural shifts in the Republic of Sakha (Yakutia) with potential relevance for past wildfire dynamics. Sources: 1: Kuzmin, 2023; 2: Kuzmin et al., 2022; 3: Crate (2021); 4: Solovyeva et al. (2022); 5: Vinokurova et al. (2022); 6: Pyne (1996); 7: Takakura et al. (2021).

Human activity intensified at the onset of the Bronze Age (c. 5000 years BP; Fig. 5), coinciding with a shift in the fire-climate relationship (Fig. 3C). The number of people following south-to-north migration towards the Siberian Arctic increased strongly (Kuzmin, 2023, Sikora et al., 2019). Increased mobility may have been facilitated by the proliferation of horse riding (Kuzmin et al., 2022). Central Yakutia, connecting the Baikal region to the Arctic via the Lena River, was an important region for major migrations (Kuzmin, 2023). In addition, animal husbandry and agriculture began spreading from Central and East Asia towards southern Siberia and the Far East around that time (Kuzmin et al., 2022), mixing with or displacing hunter-gatherer communities (Nomokonova et al., 2010). Nomadic pastoralism with sheep, goats, cattle, and horses has been recorded in the Cis-Baikal region, south of Yakutia, since at least 3000 years BP (Nomokonova et al., 2010). How early these developments spread northwards into Yakutia remains difficult to assess, but based on our results, we suggest that there may have already been impacts on wildfire dynamics during the Bronze Age. Low population density does not exclude the possibility of human impacts on fire regimes by itself (Dietze et al., 2018; Roos et al., 2018). The spread of pastoralist livelihoods in the Late Holocene may have sufficiently reduced fuel loads to alter regional fire regimes (Rouet-Leduc et al., 2021), especially along the main migration routes. Considering increased population growth during warmer, i.e., more fire-prone climatic periods (Yu et al., 2023), this could also explain the negative correlation between reconstructed biomass burning and the climate-driven simulations during the Late Holocene.

The fit between our simulated and reconstructed wildfire activity improves when accounting for human-caused fuel reduction in the forest model (Fig. 4). In addition to wildfire dynamics that are not explained by climatic or vegetation changes alone, these factors present first evidence for the potential of land management impacts of early cultures on fire regimes in eastern Siberia during the Late Holocene. As an example, we focus on a known cultural shift around 1200 CE (1100–1300 CE), a warm climatic period (Mann et al., 2009), when the pastoralist Sakha people settled in Central Yakutia (Fig. 5; Okladnikov, 1970; Zlojutro et al., 2009; Fedorova et al., 2013; Keyser et al., 2015). Coming from southern steppe environments, the Sakha likely initially settled along the Lena River (Okladnikov, 1970). The region between the Lena, Aldan, and Amga rivers later recorded the highest population in the 17^th^ century and most of our studied lakes are located there (Dolgikh, 1960; Pakendorf et al., 2006; Fig. 1). The Sakha followed semi-nomadic to sedentary pastoralist livelihoods with horse and cattle breeding (Okladnikov, 1970; Pakendorf et al., 2006). Haymaking was an important practice to feed the animals during Yakutia’s long winters. Surrounded by dense forest, they used meadows in non-forested permafrost drained lake basins (i.e., alaas landscapes; Soloviev, 1959; Crate et al., 2017; Baisheva et al., 2023) during the summers and kept them suitable for haymaking by removing shrubs or encroaching trees (Crate, 2021). They also cleared space in the forests to create open areas for haymaking, and later for growing wheat (Naumov et al., 2020), which included the managed use of fire (Crate, 2021; Solovyeva et al., 2022; Vinokurova et al., 2022). Horses were grazing freely in forests and pastures. Pine and larch wood served as preferred construction material for houses, sheds, and other structures (Okladnikov, 1970). Sakha families did not reside permanently in consolidated settlements, but also lived by their dispersed alaases and managed their surrounding environment during the warm season (Crate, 2021). These combined practices may have succeeded in keeping severe wildfires at a distance by reducing fuel loads in the surroundings of the dispersed settlements (Vinokurova et al., 2022; Girardin et al., 2024). Macroscopic charcoal as a proxy of local to regional biomass burning emphasizes impacts of human activity in the vicinity of lake systems (Whitlock and Larsen, 2001). Testing this proposed human impact by including an artificial reduction of fuel availability after 1200 CE manages to improve the fit of simulated burned area to reconstructed biomass burning (Fig. 4).

Our results display an increase of biomass burning over the most recent centuries (Fig. 2A), which may reflect another cultural shift after the colonization of Yakutia by the Tsardom of Russia in the 17^th^ century (Okladnikov, 1970; Keyser et al., 2015; Glückler et al., 2021; Fig. 5). A rapidly growing population and expanded land development across Yakutia may have increased the rate of unintentional anthropogenic ignitions (Achard et al., 2007; Shvetsov et al., 2021). By the 20^th^ century professional fire suppression was established, while slash-and-burn agriculture was restrained (Pyne, 1996). During the collectivization period, settlements were consolidated and kolkhozes worked to increase agricultural output (Crate, 2021). Cultural burning practices were completely prohibited in Yakutia in 2015, despite opposition from Sakha locals (Solovyeva et al., 2022; Vinokurova et al., 2022). Previously managed fields further away from the newly formed towns were abandoned after the end of Soviet era collectivization (Takakura et al., 2021), which may further contribute to a build-up of fuel. These recent developments highlight the potential value of understanding how Indigenous communities may have reduced wildfire severity around their settlements throughout the past millennia.

### Conclusions

We were able to determine imprints of climate, vegetation, and humans on reconstructed wildfire dynamics by applying a combination of paleoecological and modeling approaches. Our results strengthen the understanding of Holocene wildfire activity in the unique ecosystem of eastern Siberia, which was previously underrepresented in global paleofire compositions. We find that climate was likely the main driver of longer, millennial-scale trends of wildfire activity, which implies a continuing sensitivity of wildfire regimes to climate change. A negative correlation between our simulated climate-driven burned area and reconstructed biomass burning, together with a stable vegetation composition, point towards human activity as a potential driver of shorter, centennial-scale variability of wildfire activity since c. 5000 years BP. Increasing human mobility and the spread of pastoralist livelihoods may have reduced fuel availability among major migration routes, whereas the pastoralist Sakha people later used cultural burning to shape the surrounding forest to their needs. A corresponding artificial fuel reduction in our forest model results in an improved fit of simulated to reconstructed wildfire activity during the last millennium, which further demonstrates that wildfire dynamics may have been influenced by humans earlier than expected. This finding may indicate that historical low-severity fire regimes may to some extent be a result of human landscape modification. Considering the lack of systematic assessments of past human influence on fire regimes in boreal eastern Siberia, we recommend including the evaluation of a human dimension in future paleoecological studies and to take into account traditional ecological knowledge and potential effects of past land management practices.

## Methods

### Location

The Republic of Sakha (Yakutia), located in eastern Siberia, is Russia’s largest administrative subdivision. The region is characterized by vast boreal forest dominated by *Larix* on deep, continuous permafrost (Fedorov et al., 2018). Degradation of ice-rich permafrost can lead to the development of thermokarst lakes and, over longer timescales, non-forested basins with residual lakes (alaas; Baisheva et al., 2023). The extremely continental climate can reach an annual temperature amplitude of more than 100°C between the warmest and coldest day (Glückler et al., 2022). However, despite the extremely cold winters, warm summers and relatively low annual precipitation of c. 200–400 mm create optimal conditions for frequent, but generally low-intensity surface fires (Kharuk et al., 2021), although recent years have seen increases in both fire extent and severity (Köster et al., 2021).

The studied lakes are located on a c. 700 km long transect between the republic’s capital Yakutsk in the west and Oymyakon in the east, with the westernmost site being Lake 437 (N 62.34470; E 130.37543) near the Lena river, and the easternmost site being Lake 410 (N 63.23035; E 142.95733) in the Oymyakon Highlands, and share a strongly continental climate and a similar vegetation composition. Since it was not possible to find out the given names of all visited lakes, they will be here referred to instead by their fieldwork ID (i.e., Lake 402 to Lake 455). Known lake names, as well as a compilation of general attributes of each site, are included in Supplement 1.

### Fieldwork and sediment core subsampling

Fieldwork took place in August and September 2021 (Glückler et al., 2023). A total of 66 lakes of various settings were visited, from 58 of which sediment cores were obtained. With the help of point measurements using Hondex PS-7 ultrasonic depth sounders the deepest parts of the lakes were estimated. A UWITEC gravity corer (UWITEC GmbH, Austria) was used for sediment coring from rubber boats, occasionally equipped with a hammer module. Most sediment cores were obtained in PVC tubes of 9 cm in diameter, although for a few lakes an alternative setup with 6 cm diameter was used. Sediment cores were tightly sealed at the campsites. Of all the sediment cores presented in this study, only the one from Lake 408 (sediment core EN21408-2) was subsampled in the field in consecutive 1 cm intervals and stored in Whirl-Pak® bags. After the expedition, all sediment cores were collected at the North-Eastern Federal University in Yakutsk before being shipped to Potsdam, Germany, in cooled thermoboxes, where they were stored at 4°C.

In April 2022 one sediment core from each lake was selected based on length and visual quality, and opened lengthwise with an electric saw in a clean climate chamber. The sediment core inside the opened tube was split into two halves using long metal sheets. One half of each sediment core was designated for subsampling, with the other half being archived. From all 58 available sediment cores, 14 were selected for subsampling based on lake location, ensuring coverage along the whole expedition route, as well as length, diameter, and a visual confirmation of undisturbed sedimentation. Between July and November 2022, these 14 sediment cores were subsampled in contiguous 1 cm segments in a clean climate chamber under sterile conditions. To avoid contamination the topmost sediment surface and all sides touching the PVC tube were carefully removed using sterile scalpels. From each segment we obtained samples for charcoal and palynological analysis (1 cm^3^) and for total organic carbon measurement (TOC; 1–2 cm^3^), with excess sediment material being stored for potential other analyses. Every 10 cm we extracted a bulk sediment sample for radiocarbon (^14^C) dating (2 cm^3^). For the sediment core from Lake 408, subsampled in the field, the same amounts of sediment were taken out of the individual Whirl-Pak® bags for charcoal and palynological analysis, and ^14^C dating, respectively.

### ^14^C dating

Bulk sediment ^14^C samples were freeze-dried and homogenized in a planetary mill. TOC was measured with a soliTOC cube analyzer. Accelerator mass spectrometry ^14^C dates were obtained at the MICADAS (Mini Carbon Dating System) laboratory at AWI Bremerhaven, Germany, following standard protocols (Mollenhauer et al., 2021; Supplement 2, 3). ^14^C ages were calibrated using the IntCal20 calibration curve (Reimer et al., 2020) during age-depth-modeling with the package “rbacon” (Blaauw and Christen, 2011) in R (v. 4.3.2; R Core Team, 2023). Based on the resulting sediment core chronologies, another selection was made to include only those with valid age-depth relationships, resulting in the eight cores included in this study. Following Walker et al. (2012), the periods of Early, Mid-, and Late Holocene refer to subdivisions at 8200 and 4200 years BP, respectively.

### Macroscopic charcoal analysis

Preparation of macroscopic charcoal samples was done by the well-established wet sieving approach, following previously published protocols (Glückler et al., 2021, 2022). Sediment samples were soaked in a solution of sodium pyrophosphate (Na_4_P_2_O_7_) for 1–3 days to ease disaggregation of the sediment matrix. To enable the determination of pollen concentrations, one tablet of *Lycopodium clavatum* marker spores (Lund University, Department of Geology) per sample was dissolved in 10% hydrochloric acid (HCl) and added. Then, the samples were poured into a sieve at 150 µm mesh width, a standard size to separate macroscopic charcoal from the smaller sediment and pollen fractions. After thorough sieving, the macroscopic fraction in the sieve was transferred into 50 mL falcon tubes. After letting the samples rest, they were carefully decanted. Next, c. 20 mL of sodium hypochlorite (NaClO) bleach were added and samples left to soak overnight. This bleaching step improves charcoal identification by increasing the contrast between the black, non-reactive charcoal particles against other bleached organic matter (Hawthorne et al., 2018). In a final step, samples were briefly rinsed in a 63 µm mesh sieve to improve clarity after bleaching.

Charcoal quantification was done in a gridded petri dish under a Zeiss Stemi SV 11 stereomicroscope. All charcoal particles per sample were counted and grouped according to size classes and morphology. Size classes (“small”: 150–300 µm, “medium”: >300–500 µm, “large”: >500 µm) were estimated by measuring a particle’s longest axis with needles of a known tip diameter (Glückler et al., 2022). For charcoal morphology we applied the classification of morphotypes by Enache and Cumming (2007), and further grouped these into “irregular”, “angular”, and “elongated” morphologies (Glückler et al., 2022). For four of the sediment cores the length to width (L:W) ratios of all counted particles were recorded, determined with the help of an ocular micrometer (Supplement 4).

Charcoal concentrations with age information were interpolated to the median temporal resolution of each sediment core, before calculating charcoal accumulation rates (CHAR), using the “pretreatment” function of the R package “paleofire” (v1.2.4; Blarquez et al., 2014; Supplement 5). CHAR represents wildfire activity as the amount of biomass burned, integrating individual fire regime attributes of extent, intensity, or severity (Adolf et al., 2018; Hennebelle et al., 2020). Using the same R package, a charcoal composite curve was created, following Blarquez et al. (2014). A total of 12 charcoal records was used, including the eight new charcoal records presented in this study, as well as four records previously published and available in the Global Paleofire Database (GPD; Power et al., 2010; available at www.paleofire.org; data obtained in April 2024): Lake Sugun (Katamura et al., 2009a), Maralay Alaas (Katamura et al., 2009b), Lake Khamra (Glückler et al., 2021), and Lake Satagay (Glückler et al., 2022). Other charcoal records in Yakutia registered in the GPD were excluded, either because they only recorded microscopic charcoal on non-contiguous pollen slide samples, and/or because they lacked a contiguous sampling scheme for macroscopic charcoal. All charcoal records were manually added to the “paleofire” R package routine in the standard “CharAnalysis” file format (Higuera et al., 2009; function “pfAddData”). Following common recommendations (Blarquez et al., 2014; Power et al., 2008), charcoal records were transformed using a MinMax re-scaling, a Box-Cox transformation for homogenizing variance across records (Box and Cox, 1964), and a Z-score standardization, all for the base period of -70 to 11,000 years BP (function “pfTransform”). The composite curve was created following an established method (Marlon et al., 2009; Daniau et al., 2012), binning individual records into non-overlapping 100-year bins and smoothing each binned record using a locally weighted scatterplot smoothing (LOWESS) at a window width of 1000 years (function “pfCompositeLF”). Confidence intervals were generated using the distribution of 1000 bootstrapped replicates from the binned records (Blarquez et al., 2014).

### Simulations in LAVESI-FIRE

We applied the individual-based, spatially explicit forest model LAVESI (*Larix* Vegetation Simulator), which has been described in detail in previous publications (Kruse et al., 2016, 2018, 2022). Specifically, we used the fire-enabled version LAVESI-FIRE as presented by Glückler et al. (2024). In short, a simulation in LAVESI-FIRE consists of individual trees and seeds and a litter layer on top of a permafrost active layer, all within a gridded simulation area.

The model simulates annual cycles of individual tree establishment, growth, competition, and mortality, driven by climatic forcing data of monthly temperature and precipitation, as well as underlying environmental data (elevation, slope, topographic wetness index). It is capable of representing climate-driven wildfire activity, where, based on a monthly determined fire probability, fires can stochastically occur within the simulation area and impact tree and seed mortality and the litter layer height, depending on local fire intensity. To represent fuel-fire relationships, we here additionally implemented the possibility of changing the fuel availability (represented by local tree density and litter layer height) to mediate each affected grid cell’s fire intensity (Supplement 6).

We localized the model’s environmental inputs for an exemplary simulation area of 1980 × 1980 m, located in a representative alaas landscape between our study sites in Central Yakutia, using the TanDEM-X 30 m digital elevation model product (Rizzoli et al., 2017). For climatic forcing we used modeled data from MPI-ESM-CR spanning 25,000 years BP (Kapsch et al., 2022), localized at the simulation area location by monthly mean-fitting to the corresponding overlapping period with the interpolated observational data from CRU-TS v4.07 (Harris et al., 2020). The fire module was localized following Glückler et al. (2024), establishing a linear model between monthly burned area, represented by burned pixels from satellite observations between 2001 to 2022 CE in a 200 km buffer around Yakutsk (MCD64A1 product; Giglio et al., 2018), and temperature and precipitation from CRU-TS v4.07. Thresholds for mild, severe, and extreme monthly fire probability were set as third and fourth quantiles of the distribution of all fire probability values generated with the MPI-ESM-CR climate input.

We ran ten simulations for each of two scenarios, respectively. To test the impact of climatic forcing on fire regime changes, in the “normal fuel”-scenario no changes were made regarding the model’s fuel availability. In the “reduced fuel”-scenario, however, simulations ran at reduced fuel availability (20%) beginning in 1200 CE and to present. This was done to test whether such an artificial fuel reduction improves the fit between simulated and reconstructed trends of biomass burning. LAVESI-FIRE simulation output of annual burned area, corresponding to the number of burned grid cells of the simulation area, was smoothed with locally estimated scatterplot smoothing (LOESS; span ranging from 0.05 for multi-centennial trends to 0.5 for multi-millennial trends). Then, the median and interquartile range of all smoothed time series of the ten simulation repeats per scenario were used for visualization. Spearman rank correlation coefficients (*r*_s_) are used to assess associations between the non-normally distributed data of simulated burned area and the charcoal composite curve.

## Data availability

Charcoal and radiocarbon age data presented in this study are available via PANGAEA (https://doi.pangaea.de/10.1594/PANGAEA.974511 and https://doi.pangaea.de/10.1594/PANGAEA.974676; in review) and will be made available in the Global Paleofire Database (https://paleofire.org/).

## Code availability

The LAVESI-FIRE simulation output is available via Zenodo (https://zenodo.org/records/14626969; in review), and the model code can be accessed via GitHub (https://github.com/StefanKruse/LAVESI/tree/fire).

## Acknowledgements

We thank the team of the German-Russian “Yakutia 2021” expedition. W. Finsinger kindly provided support with R. Thanks to P. Uchanov for help in the laboratory and to everyone supporting sediment core subsampling: J. Courtin, V. Döpper, A. Eichner, L. Enguehard, L. Grimm, S. Haupt, P. Hauter, A. Korup, H.S. Malik, P. Meister, C. Messner, R. Paasch, A. Prasannakumar, K. Schildt, E. Topp-Johnson, and J. Wagner.

## Author contributions

U.H., S.K., and L.P. designed and led the fieldwork, supported by R.G. R.G., U.H., E.Z., I.B., and A.S. conducted fieldwork at the lake sites. R.G. designed the charcoal analysis study, supervised by U.H and E.D. R.G. subsampled the sediment cores. R.G. prepared the age dating process and created the chronologies. R.G. conducted the charcoal-related laboratory work and data analysis, supported by S.T. and E.D. R.G. wrote the initial version of the manuscript, supervised by U.H and supported by A.A. All authors reviewed the initial manuscript.

