## Supplement for "Human dimension of Holocene wildfire dynamics in boreal eastern Siberia"

| Location | Highlands | | | Lowlands | | | | |
|---|---|---|---|---|---|---|---|---|
| Sediment core | EN21402-4 | EN21408-2 | EN21410-1 | EN21419-2 | EN21421-2 | EN21433-1 | EN21437-1 | EN21455-2 |
| ID, this study | 402 | 408 | 410 | 419 | 421 | 433 | 437 | 455 |
| Lake name | Unknown | Unknown | Unknown | Lankyly (Ланкылы) | Nidzhili (Нидьили) | Unknown | Unknown | Unknown |
| Region | Verkhoyansk Mountains | Verkhoyansk Mountains | Oymyakon Highlands | Churapcha | Churapcha | Tungulu | Tungulu | Maya |
| Lake type | Intermontane basin lake, glacial lake | Intermontane basin lake, glacial lake | Intermontane basin lake | Thermokarst lake (alaas) | Thermokarst lake (alaas) | Thermokarst lake (alaas) | Thermokarst lake (alaas) | Thermokarst lake (alaas) with pingo (bulgunyakh) |
| Latitude<br>Longitude | 63.32696<br>141.08434 | 63.38775<br>140.57524 | 63.23035<br>142.95733 | 62.08767<br>132.35340 | 62.01409<br>132.41266 | 62.13806<br>130.87054 | 62.3433<br>130.37467 | 61.90249<br>130.41174 |
| Elevation (m a.s.l.) | 989 | 1254 | 789 | 289 | 160 | 229 | 127 | 144 |
| Water depth (m) | 12.6 | 8.9 | 7.4 | 3.4 | 1.7 | 7.0 | 0.5 | 1.0 |
| Lake area (ha) | 32.0 | 11.2 | 5.7 | 14.8 | 24.3 | 6.3 | 3.4 | 27.9 |
| Fieldwork date (YYYY-MM-DD) | 2021-08-08 | 2021-08-12 | 2021-08-13 | 2021-08-18 | 2021-08-19 | 2021-08-24 | 2021-08-25 | 2021-08-30 |
| Sediment recovered (cm) | 37.4 | 28.0 | 29.8 | 38.6 | 32.4 | 68.0 | 41.0 | 35.0 |
| Samples for this study (n) | 36 | 28 | 29 | 38 | 32 | 66 | 40 | 34 |
| Basal age (years BP) | c. 4960 | c. 2410 | c. 4740 | c. 620 | c. 1180 | c. 4410 | c. 7720 | c. 1090 |
| Mean temporal resolution per sample (years) | c 138 | c. 86 | c. 163 | c. 16 | c. 37 | c. 67 | c. 193 | c. 32 |

**Supplement 1:** Table of lake metadata.

| Lake ID | Sediment core | Lab ID | Depth top (cm) | Depth bot (cm) | F$^{14}$C | ± 1σ | $^{14}$C years BP | ± 1σ | cal. years BP | ± 2σ | Comment |
|---|---|---|---|---|---|---|---|---|---|---|---|
| 402 | EN21402-4 | 9764.1.1 | 10 | 11 | 0.79 | 0.0026 | 1858 | 26 | 1769 | 56 | |
| 402 | EN21402-4 | 9765.1.1 | 20 | 21 | 0.71 | 0.0024 | 2772 | 27 | 2810 | 27 | |
| 402 | EN21402-4 | 9766.1.1 | 35 | 36 | 0.59 | 0.0022 | 4293 | 30 | 4863 | 36 | |
| 408 | EN21408-2 | 10214.1.3 | 10 | 11 | 0.82 | 0.0022 | 1590 | 22 | 1467 | 58 | |
| 408 | EN21408-2 | 10215.1.3 | 20 | 21 | 0.77 | 0.0021 | 2124 | 22 | 2021 | 19 | |
| 408 | EN21408-2 | 10216.1.3 | 27 | 28 | 0.76 | 0.0021 | 2234 | 22 | 2225.5 | 71.5 | |
| 410 | EN21410-1 | 9767.1.1 | 10 | 11 | 0.88 | 0.0027 | 1046 | 25 | 976 | 58 | |
| 410 | EN21410-1 | 9768.1.1 | 28 | 29 | 0.59 | 0.0018 | 4174 | 24 | 4602.5 | 14.5 | |
| 419 | EN21419-2 | 9769.1.1 | 10 | 11 | 1.12 | 0.0032 | | | | | Excluded from age-depth modeling |
| 419 | EN21419-2 | 9770.1.1 | 20 | 21 | 0.95 | 0.0029 | 374 | 24 | 353.5 | 33.5 | |
| 419 | EN21419-2 | 9771.1.1 | 37 | 38 | 0.93 | 0.0028 | 606 | 24 | 565.5 | 17.5 | |
| 421 | EN21421-2 | 10220.1.2 | 10 | 11 | 0.97 | 0.0026 | 252 | 21 | 77 | 77 | |
| 421 | EN21421-2 | 10221.1.2 | 20 | 21 | 0.88 | 0.0024 | 1023 | 22 | 935.5 | 22.5 | |
| 421 | EN21421-2 | 10222.1.2 | 31 | 32 | 0.87 | 0.0023 | 1163 | 22 | 1013 | 37 | |
| 433 | EN21433-1 | 9774.1.1 | 10 | 11 | 0.43 | 0.0019 | 6790 | 36 | 7628.5 | 49.5 | Considered outlier by age-depth modeling in "rbacon" |
| 433 | EN21433-1 | 9775.1.1 | 30 | 31 | 0.86 | 0.0024 | 1212 | 22 | 1120 | 54 | |
| 433 | EN21433-1 | 9776.1.1 | 50 | 51 | 0.69 | 0.0020 | 3031 | 23 | 3225.5 | 61.5 | |
| 433 | EN21433-1 | 9777.1.1 | 65 | 66 | 0.61 | 0.0026 | 3946 | 35 | 4271.5 | 18.5 | |
| 437 | EN21437-1 | 10223.1.2 | 10 | 11 | 0.77 | 0.0021 | 2106 | 22 | 2068 | 70 | |
| 437 | EN21437-1 | 10224.1.2 | 20 | 21 | 0.60 | 0.0017 | 4071 | 23 | 4478.5 | 35.5 | |
| 437 | EN21437-1 | 10225.1.2 | 30 | 31 | 0.52 | 0.0015 | 5197 | 24 | 5953 | 41 | |
| 437 | EN21437-1 | 10226.1.2 | 39 | 40 | 0.44 | 0.0013 | 6549 | 24 | 7485.5 | 62.5 | |
| 455 | EN21455-2 | 10231.1.2 | 10 | 11 | 0.90 | 0.0024 | 863 | 22 | 710 | 13 | |
| 455 | EN21455-2 | 10232.1.2 | 20 | 21 | 0.90 | 0.0024 | 870 | 22 | 801.5 | 73.5 | |
| 455 | EN21455-2 | 10233.1.1 | 33 | 34 | 0.88 | 0.0024 | 1054 | 22 | 954 | 32 | |

**Supplement 2:** Table of sediment core radiocarbon age dating information. All radiocarbon data can be accessed via PANGAEA (see main manuscript).

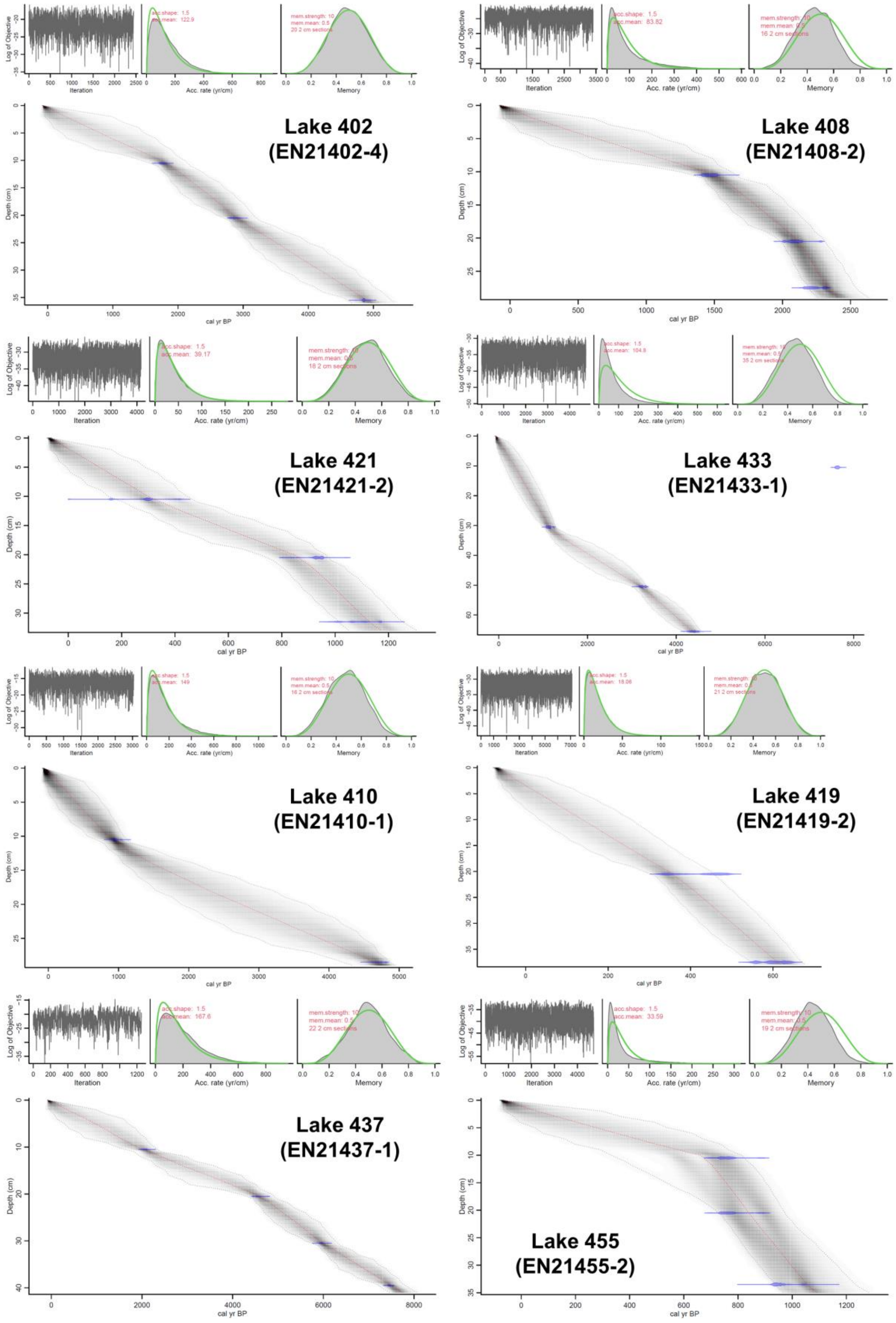


**Supplement 3:** Individual sediment core chronologies. All radiocarbon data can be accessed via PANGAEA (see main manuscript).

| Lake ID | Mean charcoal conc. (# cm$^{-3}$) | Mean CHAR (# cm$^{-2}$ yr$^{-1}$) | Small size (%) | Medium size (%) | Large size (%) | Angular types (%) | Elongated types (%) | Irregular types (%) | Mean L:W |
|---|---|---|---|---|---|---|---|---|---|
| 402 | 6.3 ± 5.2 | 0.04 ± 0.04 | 40.7 | 34.1 | 25.2 | 29.6 | 16.4 | 54.0 | 4.13 |
| 408 | 3.3 ± 2.2 | 0.04 ± 0.04 | 58.7 | 26.1 | 15.2 | 38.0 | 12.0 | 50.0 | - |
| 410 | 20.4 ± 21.9 | 0.13 ± 0.12 | 33.6 | 27.0 | 39.4 | 61.1 | 24.2 | 14.7 | 5.41 |
| 419 | 2.6 ± 1.8 | 0.14 ± 0.09 | 55.6 | 26.3 | 18.2 | 28.3 | 31.3 | 40.4 | 3.46 |
| 421 | 50.3 ± 27.1 | 1.27 ± 1.08 | 45.1 | 27.0 | 27.9 | 63.4 | 5.5 | 31.2 | 2.90 |
| 433 | 14.6 ± 13.6 | 0.21 ± 0.18 | 55.5 | 28.3 | 16.3 | 53.5 | 18.0 | 28.5 | - |
| 437 | 57.8 ± 24.7 | 0.29 ± 0.11 | 62.7 | 27.2 | 10.2 | 72.3 | 11.2 | 16.6 | - |
| 455 | 17.8 ± 6.2 | 0.52 ± 0.49 | 57.5 | 29.5 | 13.1 | 78.0 | 12.4 | 9.6 | - |

**Supplement 4:** Table with additional charcoal record data for the eight new charcoal records.

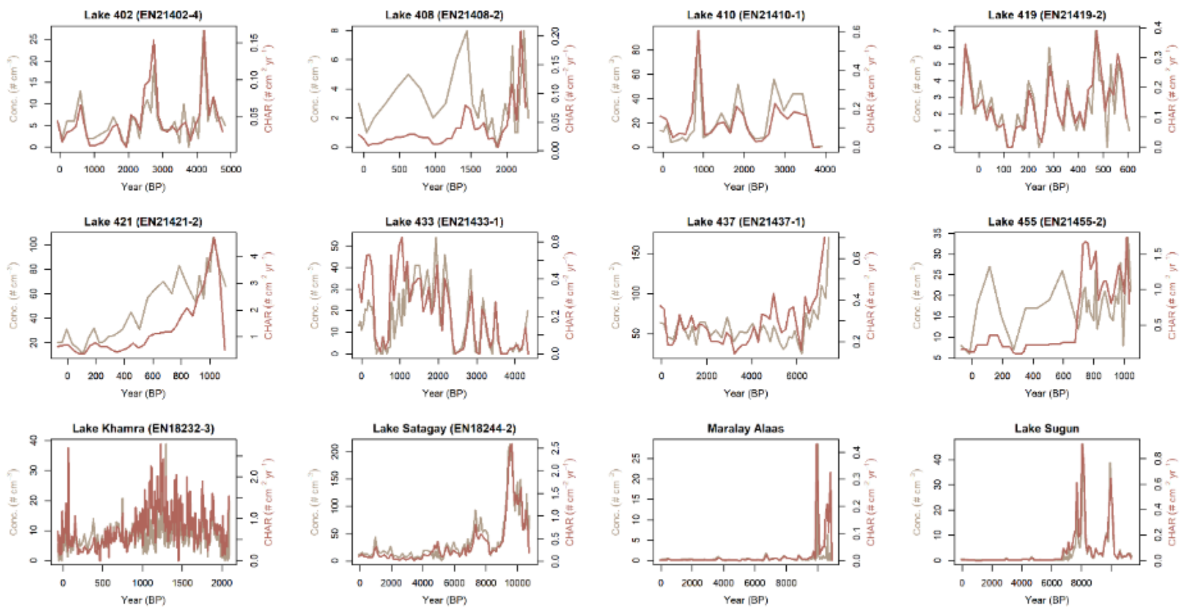


**Supplement 5:** Individual charcoal records included in the composite for the Republic of Sakha (Yakutia), showing both concentration values and charcoal accumulation rates (CHAR). Lake 402-455: New contributions of this study. Lake Khamra: Glückler et al. (2021). Lake Satagay: Glückler et al. (2022). Maralay Alaas: Katamura et al. (2009b). Lake Sugun: Katamura et al. (2009a). Full references can be found in the main manuscript.

$$FF = \left(-1 * \left(\frac{LH}{3000}\right) * FA\right) + \left(\frac{TD}{65535}\right) * FA$$

FF: Fuel factor
LH: Litter layer height (0-3000; 3000 equals 30 cm)
FA: Fuel availability (0-1; custom parameter that defaults to 1)
TD: Tree density (0-65535)

**Supplement 6:** Fuel availability implementation in LAVESI-FIRE (Glückler et al., 2024). Full reference can be found in the main manuscript.
